# Fatal toxicity of chloroquine or hydroxychloroquine with metformin in mice

**DOI:** 10.1101/2020.03.31.018556

**Authors:** N.V Rajeshkumar, Shinichi Yabuuchi, Shweta G. Pai, Anirban Maitra, Manuel Hidalgo, Chi V. Dang

**Affiliations:** Department of Oncology, Pathology, Radiation Oncology and Molecular Radiation Sciences, Johns Hopkins University School of Medicine, Baltimore, MD; Department of Pathology, Johns Hopkins University School of Medicine, Baltimore, MD; Sheikh Ahmed Center for Pancreatic Cancer Research, M.D. Anderson Cancer Center, Houston, TX; Division of Hematology and Medical Oncology, Weill Department of Medicine, Sandra and Edward Meyer Cancer Center, Weill Cornell Medicine, New York, NY; Ludwig Institute for Cancer Research, New York, NY

## Abstract

Guided by the principle of *primum non nocere* (first do no harm), we report a cautionary note on the potential fatal toxicity of chloroquine (CQ) or hydroxychloroquine (HCQ) in combination with anti-diabetic drug metformin. We observed that the combination of CQ or HCQ and metformin, which were used in our studies as potential anti-cancer drugs, killed 30-40% of mice. While our observations in mice may not translate to toxicity in humans, the reports that CQ or HCQ has anti-COVID-19 activity, the use of CQ resulting in toxicity and at least one death, and the recent Emergency Use Authorization (EUA) for CQ and HCQ by the US Food and Drug Administration (FDA) prompted our report. Here we report the lethality of CQ or HCQ in combination with metformin as a warning of its potential serious clinical toxicity. We hope that our report will be helpful to stimulate pharmacovigilance and monitoring of adverse drug reactions with the use of CQ or HCQ, particularly in combination with metformin.

## Introduction

Guided by the principle of *primum non nocere* (first do no harm), we report a cautionary note on the potential fatal toxicity of chloroquine (CQ) or hydroxychloroquine (HCQ) in combination with anti-diabetic drug metformin. We observed that the combination of CQ or HCQ and metformin, which were used in our studies as potential anti-cancer drugs, killed 30-40% of mice. While our observations in mice may not translate to toxicity in humans, the reports that CQ or HCQ has anti-COVID-19 activity [1], the use of CQ resulting in toxicity and at least one death, and the recent Emergency Use Authorization (EUA) for CQ and HCQ by the US Food and Drug Administration (FDA) prompted our report. Here we report the lethality of CQ or HCQ in combination with metformin as a warning of its potential serious clinical toxicity. We hope that our report will be helpful to stimulate pharmacovigilance and monitoring of adverse drug reactions with the use of CQ or HCQ, particularly with metformin.

## Methods

Animal experiments were conducted following approval by the Animal Care and Use Committee guidelines of the Johns Hopkins University (Baltimore, MD). Tumor bearing or non-tumor bearing immunocompromised mice were injected with 100 μL of saline vehicle, chloroquine (CQ, 60 mg/kg), hydroxychloroquine (HCQ, 60 mg/kg) and/or metformin (250 mg/kg) once daily intraperitoneally daily for 4 weeks as described [2 3]. A combination of CQ and metformin in the above-mentioned dose and frequency was administered to animals in the combination treatment group. In a separate study, non-tumor bearing immunocompromised and immunocompetent mice were treated with HCQ and metformin once daily in the above-mentioned dose for 38 days. Blood was drawn for chemistry and hematology via cardiac puncture, and organs and tissues were harvested, examined and processed for transmission electron microscopy as described [3].

## Results

Based on our previous findings that metformin and CQ or HCQ curbed the growth of human pancreatic xenografts in athymic nude mice [2 3], we sought to determine whether the combination of metformin, which inhibits mitochondrial Complex I, and CQ that inhibits autophagy could be synergistic as an anti-cancer metabolic cocktail. In contrast to single agent metformin or CQ, which have anti-tumor activity, we found that the combination of metformin and CQ was lethal in 40% of tumor bearing or non-tumor bearing mice (**Figure 1A**).

**Figure 1.**
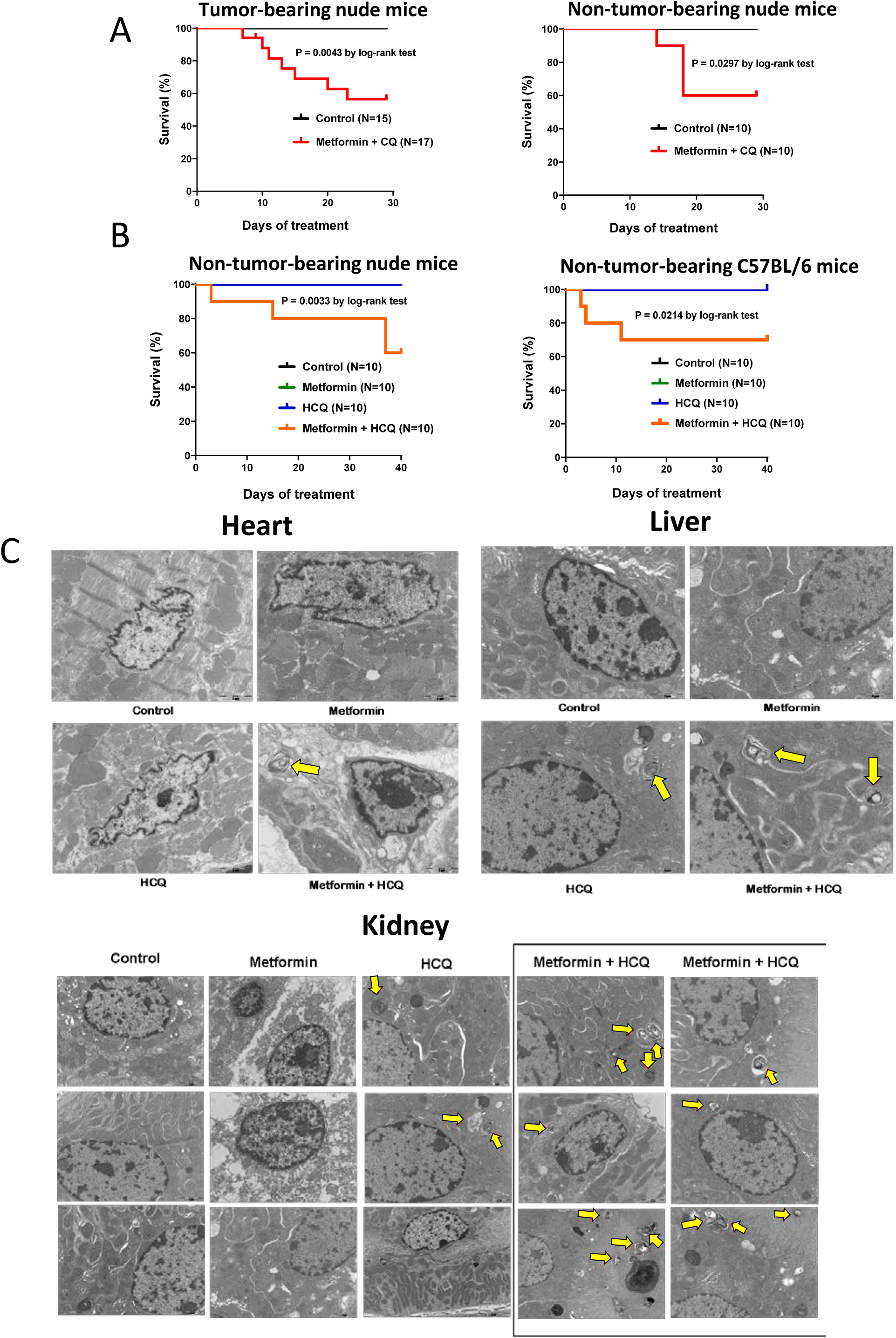

To determine whether HCQ is similarly lethal in combination with metformin, we tested the metformin+HCQ combination in non-tumor bearing nude mice, which showed a 40% mortality (**Figure 1B**). To determine whether immunocompromised nude mice were particularly sensitive to the combination, we treated immunocompetent C57BL/6 mice with the metformin+HCQ, which resulted in 30% lethality.

We then sought to determine the basis for the toxicity of metformin+HCQ combination in autopsy studies and found that body weights were not significantly different (**Table 1**) among the different groups. While organ weights were not different among the groups, we observed via transmission electron microscopy an increase in the number of autophagosomes in the heart, liver and kidneys of athymic nude mice treated with metformin+HCQ combination (**Figure 1C**). While the hematological findings were not different among the groups, we found that lactate dehydrogenase (LDH) and creatine kinase (CK) levels were elevated in all treatment groups as compared to control vehicle treated group (**Table 1**).

**Table 1.**
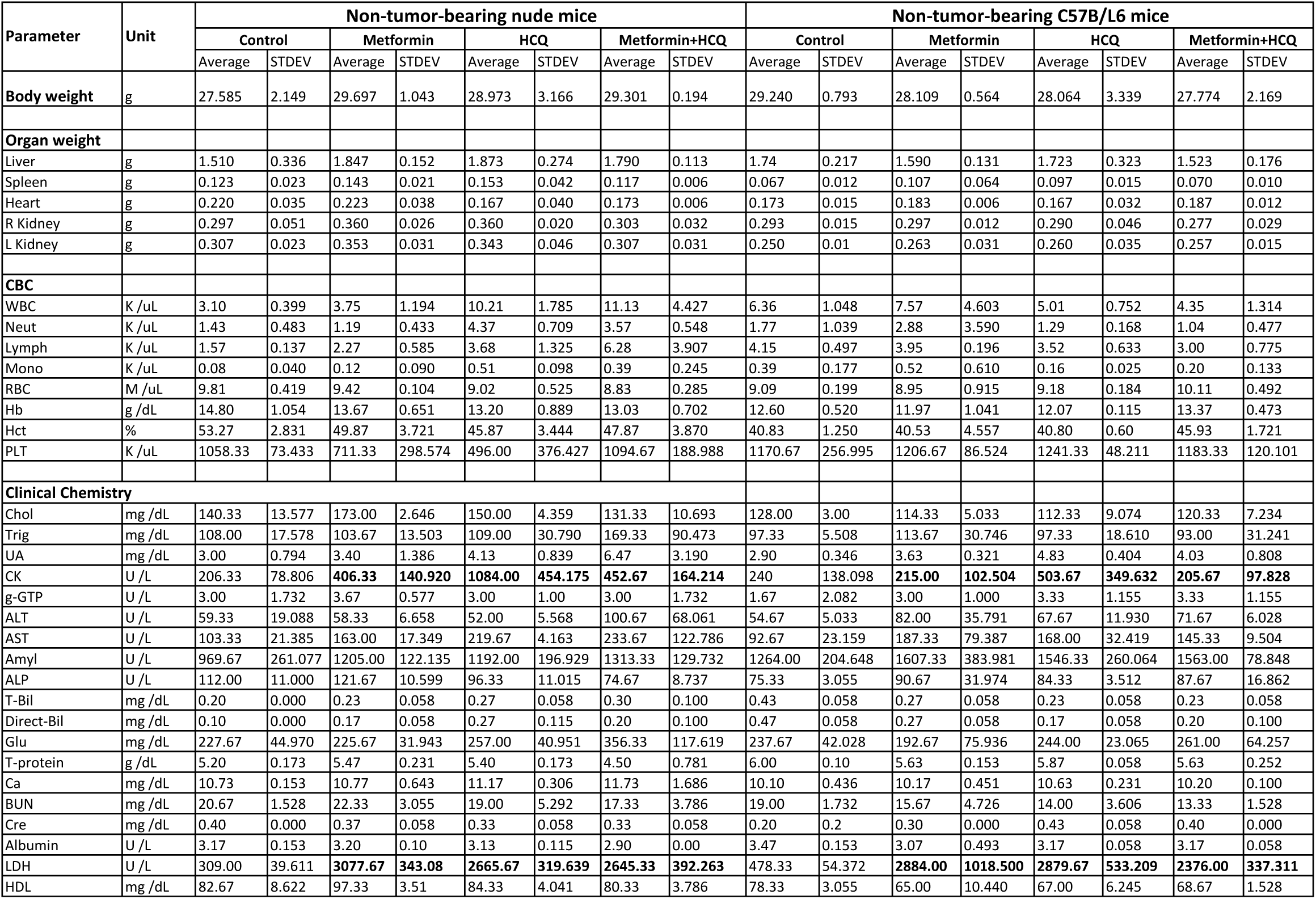
Body weight, organ weight, CBC and clinical chemistry parameters of mice administered with Vehicle (Control), Metformin, HCQ or Metformin and HCQ.

## Discussion

Between March 16 and 21, 2020, partly based on a non-randomized study using HCQ with azithromycin [4], claims disseminated through Twitter and amplified by the media that CQ or HCQ could be a therapy for COVID-19. Many individuals have started to take this drug, resulting in chloroquine poisoning in Nigeria (@NCDCgov #COVID19Nigeria) and a death in Arizona. Notably, CQ and HCQ doses which are used for the treatment of rheumatic diseases, could lead to the development of hypoglycemia, cardiomyopathy and retinopathy [5]. Here we report the lethality of metformin+CQ or +HCQ as a warning of its potential deadly toxicity, noting that the dosages in mice are similar to those in human with allometric scaling. Consistent with our findings, the combination of CQ and metformin resulted in CNS neuronal damage after cardiac arrest in rats [6]. We hope that our report will be helpful to stimulate pharmacovigilance and monitoring of adverse drug reactions with the use of CQ or HCQ, particularly in combination with metformin.

## Author contributions

*Concept and design:* Rajeshkumar, Maitra, Hidalgo, Dang

*Acquisition, analysis or interpretation of data*: Rajeshkumar, Yabuuchi, Pai, Dang, Hidalgo

*Drafting of manuscript*: Dang, Hidalgo, Rajeshkumar

*Statistical analysis*: Rajeshkumar, Yabuuchi

*Obtained funding*: Dang, Maitra, Hidalgo, Rajeshkumar

Drs. Dang and Hidalgo have access to primary data in Dr. Rajeshkumar’s possession.

